# Neural correlates of state transitions elicited by a chemosensory danger cue

**DOI:** 10.1101/2020.04.23.054734

**Authors:** Suresh Jesuthasan, Seetha Krishnan, Ruey-Kuang Cheng, Ajay Mathuru

## Abstract

**Background:** Detection of predator cues changes the brain state in prey species and helps them avoid danger. Dysfunctionality in changing the central state appropriately in stressful situations is proposed to be an underlying cause of multiple psychiatric disorders in humans.

**Methods:** Here, we investigate the dynamics of neural circuits mediating response to a threat, to characterize these states and to identify potential control networks. We use resonant scanning 2-photon microscopy for *in vivo* brain-wide imaging and custom designed behavioral assays for the study.

**Results:** We first show that 5-7 day old zebrafish larvae react to an alarm pheromone (*Schreckstoff*) with reduced mobility. They subsequently display heightened vigilance, as evidenced by increased dark avoidance. Calcium imaging indicates that exposure to *Schreckstoff* elicits stimulus-locked activity in olfactory sensory neurons innervating a lateral glomerulus and in telencephalic regions including the putative medial amygdala and entopeduncular nucleus. Sustained activity outlasting the stimulus delivery was detected in regions regulating neuromodulator release, including the lateral habenula, posterior tuberculum, superior raphe, and locus coeruleus.

**Conclusion:** We propose that these latter regions contribute to the network that defines the “threatened” state, while neurons with transient activity serve as the trigger. Our study highlights the utility of the zebrafish larval alarm response system to examine neural circuits during stress dependent brain state transitions and to discover potential therapeutic agents when such transitions are disrupted.

## Introduction

Behavioral and physiological changes in response to a danger signal increase the chances of survival in animals. These responses occur over multiple time-scales; immediate defensive behaviors that help evade predators [7] are coupled with long-term, system-wide changes to counter risk [1]. Regions of the vertebrate brain that process and execute immediate defensive behaviors, such as freeze or flight, have been well-studied [22, 29]. The role of specific brain regions in mediating different phases of sustained response, which have been variously described as sustained fear [16], anxiety-like, or anxiety-related behaviors [4] has also been investigated [16, 50]. These studies have identified evolutionarily conserved circuits mediating the response to a threat. However, a whole-brain view of the neural dynamics mediating these responses, which is critical in objectively defining brain state and identifying potential control circuits, i.e. the trigger for change in state [35], is lacking. Furthermore, unraveling the dynamics and organization of functional brain networks during stress is thought to be a central piece of the puzzle to understanding the abnormality in neuropsychiatric conditions such as mood disorders and Schizophrenia that are viewed as “disconnectivity syndromes” [11].

Here, we investigated the use of larval zebrafish, where *in vivo* brain-wide imaging of neural activity can be conducted with relative ease [13, 23], as a model to study the dynamics of neural response to danger. As a danger cue, we turned to alarm substance (or *Schreckstoff*). This is released upon physical injury to an individual and elicit a striking change in the behavior of conspecifics [15, 24, 36, 44]. There is an immediate change in locomotion [27, 45], an increase in arousal [42], and an increase in anxiety-like behaviors that persist after the removal of the cue [31, 38]. Thus, the alarm response is a good paradigm to examine changes in central states [43].

Whether early zebrafish larvae (5-7 day old) ideally suited for whole-brain imaging show a *Shreckreaktion* has been debated. One study examined the ontogeny of the response in zebrafish in detail by quantifying behavioral parameters associated with alarm in adults. This study reported that the earliest responses can be seen only around day 42 days post fertilization [48]. This mirrors reports in fathead minnows that observable responses occur only after 48-57 days post-hatching [12]. These observations are consistent with older studies that tie the initial onset of typical adult-like alarm response with the development of shoaling behavior and mixed feeding as larvae reach a juvenile stage (in most species between 28-40 days post fertilization [19]). However, although adult-like responses may appear only later in ontogeny, it is possible that larvae are capable of sensing and responding to the alarm cue in a different manner [20, 34]. Our behavioral and imaging experiments suggest that larval zebrafish show a dynamic response to *Schreckstoff* even at this early age, characterized by an acute change in behavior followed by a change in wariness.

## Materials and Methods

### Ethics statement

All experiments were carried out under guidelines approved by the IACUC of A*STAR (number 181408).

### Experimental methods

Are described in detail in extended supplementary data.

## Results

### Quantifying larval swimming behavior in a vertical column

We used a simple assay chamber that allowed observation of larval swimming behavior in a 50 mm vertical column (Supplementary Figure 1; [32];see methods). We first characterized and identified quantifiable parameters associated with normal swimming of 5-7 dpf larvae in this type of chamber over a period of 30 minutes. As described in the past for larvae of an equivalent age [10], larvae swam in short bouts (Fig 1A, 1B). Larvae explored the entire chamber but preferred to stay in the top quarter of the chamber reaching an average depth of about 14 mm (Figure 1C, Supplementary Figure 1). The average duration of each swimming bout was 246.40 ms (95% CI [245.59 ms, 247.21 ms]) and each inter-bout interval lasted 173.11 ms (95% CI [169.05 ms, 177.18 ms]). The histogram of swimming speed plotted per second (Figure 1D) showed a bimodal distribution with one peak in the 0 mm/sec bin reflecting the distribution of time spent in the two modes, swimming, and inter-bout intervals. We examined if the swimming behavior of larvae changed as a function of the time spent in the assay chamber. We examined 10-minute windows, on either side of the intended stimulus delivery time (calling them Pre and Post; arrow Figure 1F). We also included a second 10-minute window after the first post-stimulus delivery period (10’-20’ Post) to examine delayed responses if any. No differences in the distance swam (Figure 1F) or in the inter-bout intervals (Figure 1E) could be detected. Therefore, the swimming behavior of the zebrafish larvae did not show any noticeable change in the 30 minute observation period of the experiment.

**Figure 1.**
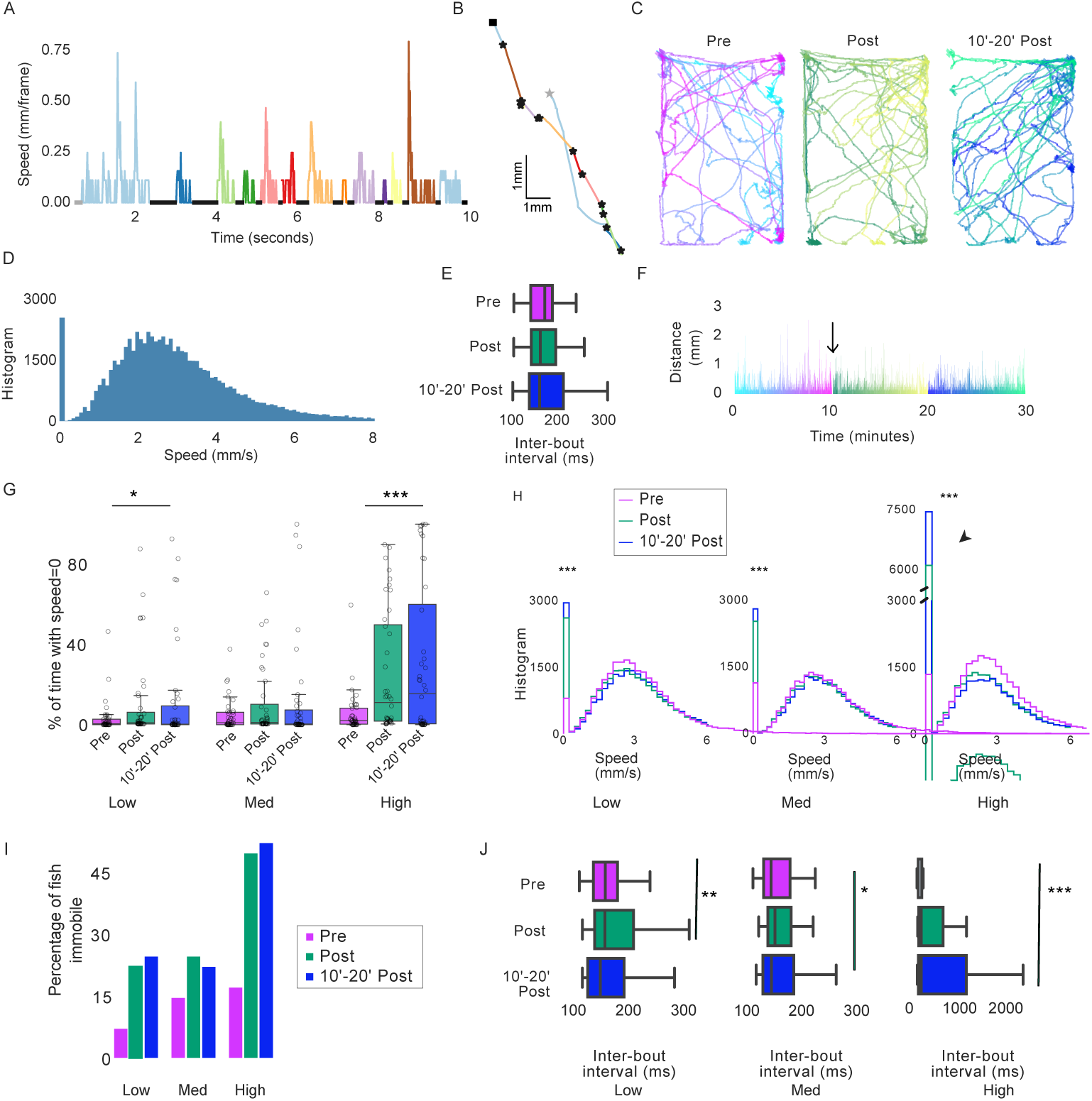
Quantification of larval behavior to *Schreckstoff* in a vertical column.. (A) Trace shows instantaneous speed over a few seconds in one larva. Swimming periods are interspersed with periods of speed = 0 (Black dots). (B) The trajectory of the same animal. Grey star indicates time = 0. Bout trajectories are color coded to panel A and black stars indicate start of a swim bout. Bouts vary in length and distance moved. Representative (C) tracks and (F) distance swam (in mm) color-coded over three 10 minute bins - Pre, Post and 10′ − 20′ Post. Stimuli, if delivered were delivered at the time indicated by the arrow in F. (D) Histogram of the speed (mm/sec) over the entire period shows the binomial distribution due to time spent in swimming bouts interspersed with inter-bout intervals of 0 speed. (E) Inter-bout intervals in the three time-bins do not change significantly over time. (G) Percentage of fish with speed = 0 mm/sec for > = 1 SD than the average immobility time in the Pre time-bin of the no stimulus condition, (H) Histograms of speed distribution (in mm/s), (I) Boxplots of the percentage of time spent motionless and (J) Inter-bout intervals in the three time-bins in three time-bins for the conditions to three concentrations of *Schreckstoff* (low, medium and high) in the Pre, Post, and 10’-20’ Post time-bins. Circles represent the individual fish response. Change is most notable in the high condition * indicate p values < 0.05, ** < 0.01, and *** indicate < 0.0001 in paired t-tests or KS test. Exact p values are given in the text; n = 40/condition

### Larval zebrafish display a startle response to stimulus delivery

Change in illumination [6], acoustic, or mechanical disturbances [33] can startle zebrafish larvae. We reasoned that a liquid stimulus delivery into the observation chamber however gently performed will disturb the water column and startle the larvae. To test this, we compared the behavior of larvae when tank water was delivered into the chamber as a control stimulus (Supplementary Figure 2). The brief stimulus delivery period (5 seconds) was excluded from the analysis. Histogram of speed distribution reflected the observations in Figure 1 that there were no differences between the three time-bins in the no stimulus delivery condition (Spplementary Figure 2).

Control (tank water) delivery however changed speed distribution (Supplementary Figure 2A). Larvae increase the time spent motionless (Supplementary Figure 2B). The change is reflected in increased inter-bout interval as the average duration of a swimming period remains unchanged at 244.99 ms (95% CI [243.58 ms, 246.40 ms]), while the average inter-bout interval increased from 171.76 ms (95% CI [168.36 ms, 175.15 ms]) to 190.89 ms (95% CI [182.06 ms, 199.71 ms]; Pre with Post, p = 0.001, Student’s t-test).

Few individuals can contribute disproportionately to an average readout, such as the one quantified above. To examine if this could be the case here, we compared the number of individuals that showed higher than average immobility in the two conditions. In the no stimulus condition, larvae spent approximately 24 seconds or 4.18 % of the 10-minute window being immobile (95% CI [1.72%, 6.64%], standard deviation = 7.60%; Supplementary Figure 2B). In the Pre time-bin, only 10% of individuals (4 of 40; Supplementary Figure 2C) were immobile for periods longer than the average by 1 SD or more (i.e. for 11.78% or longer of the 10-minute window). This percentage was unchanged in the Post and the 10’-20’Post time-bins (Supplementary Figure 2C and 2D). In the stimulus delivery condition this percentage increased to approximately 20% of individuals (9 out of 40; Supplementary Figure 2C and 2D). Therefore, a greater number of individuals were immobile for longer periods after the mechanical disturbance caused by control stimulus delivery. The inter-bout interval increased upon stimulus delivery, but the average swimming period in a bout did not change.

### Larval zebrafish respond to adult-derived *Schreckstoff*

Given that the stimulus delivery itself can elicit a detectable change in the behavior of larvae, we examined larval responses to 3 concentrations (low, medium, and high) of *Schreckstoff*. We reasoned that if larvae respond to the alarm substance then it may show a concentration dependence, while a response purely to the mechanical disturbance produced by the process of *Schreckstoff* delivery will be independent of concentration. The histogram of speed distribution showed a marked change after stimulus delivery in both low (Figure 1H; Low, Pre with 10’ Post: p=0.0002; Pre with 10’-20’ Post: p=0.00004, KS Test) and medium *Schreckstoff* conditions (Figure 1H; Med, Pre with 10’ Post: p=0.0002; Pre with 10’-20’ Post: p=0.0008, KS Test). The total time spent immobile showed a modest increase (Figure 1G). The inter-bout interval also increased after stimulus delivery from 160.38 ms (95% CI [155.52 ms, 165.23 ms]) to 198.29 ms (95% CI [180.32 ms, 216.26 ms]) in the case of low (Figure 1J; p <0.0001, Student’s t-test), and from 160.31 ms (95% CI [152.62 ms, 168.00 ms]) to 179.70 ms (95% CI [166.40 ms, 192.97 ms]) in the case of the medium concentration (Figure 1J; p = 0.012, Student’s t-test). These changes, however, were not unlike those observed for the control delivery condition described above, and a similar number of individuals (approximately 20%) showed a change in the behavior (Supplementary Figure 2).

The larval response to the highest concentration of *Schreckstoff*, on the other hand, was much more striking (Supplementary Figure 3; Supplementary Movie 1). The speed distribution histogram showed a substantial and prolonged increase in immobility (Figure 1H, SS high; Pre with 10’ Post: p=6.8*10-8; Pre with 10’-20’ Post: p=1.2*10-9, KS Test; Supplementary Figure 3). 50% of the fish (20/40) showed such a response a(Figure 1I). The duration of swimming bouts did not change much from 239.33 ms (95% CI [238.02 ms, 240.64 ms]) to 230.25 ms (95% CI [228.83 ms, 231.67 ms]), but the inter-bout interval (Figure 1J, p <0.0001, Student’s t-test) increased from 188.53 ms (95% CI [182.85 ms, 194.22 ms]) to 320.58 ms (95% CI [273.76 ms, 367.40 ms]).

Adult zebrafish show a diving response to *Schreckstoff* [37]. Supplementary Figure 5 shows the location and duration of the immobility of all the animals tested after exposure to the control stimulus (tank water) or to the highest concentration of *Schreckstoff*. The diameter of the circles is proportional to the duration of immobility as a percentage of that time-bin (10 minutes). Larvae become immobile in different parts of the chamber. Therefore, larvae show a qualitatively similar response to control stimulus delivery at low concentrations of *Schreckstoff*, but quantitatively different response when a high concentration was delivered. Greater number of fish show an immobility response. Therefore, 5-7 dpf larvae can sense and respond to adult-derived *Schreckstoff*.

### Larval zebrafish behavior changes after exposure to the alarm substance

Adult zebrafish show an increase in anxiety-like behaviors after exposure to the alarm substance, as observable by a change in their responses in the light/dark assay [38]. In such a novel environment adult zebrafish display scototaxis, that is, a preference for the dark side of the chamber. This natural response is further enhanced after exposure to *Schreckstoff* [38]. Zebrafish larvae, on the other hand, are known to display scotophobia or dark avoidance in such an assay [46].

To test whether scotophobia in larvae is also increased, indicative of lasting anxiogenic effects of transient exposure to *Schreckstoff*, we placed them in assay tanks that offered a choice between light and dark backgrounds after washing off the alarm substance. 7 dpf larvae showed an increase in scotophobia compared to control larvae exposed only to the tank water spending less time in the dark side of the tank (Figure 2A; mean difference = -5.49 [95CI = -9.39, -1.36], Cohen’s *d* = 0.6 Student’s t-test p = 0.01) and making fewer entries into the dark side (Figure 2B; mean difference = -0.9 [95CI = -0.43, -1.33], Cohen’s *d* = 0.95 Student’s t-test p = 0.0001). Alarmed larvae (n=40) are more risk averse and are more cautious compared to controls as they enter the dark side later. This is also reflected at a population level by the shallow slope of a quadratic function fit for the time to first entry to dark (Figure 2C). Therefore, zebrafish larvae exhibit anxiety-like behaviors after transient exposure to an alarm cue and remain cautious indicative of a changed central brain state.

**Figure 2.**
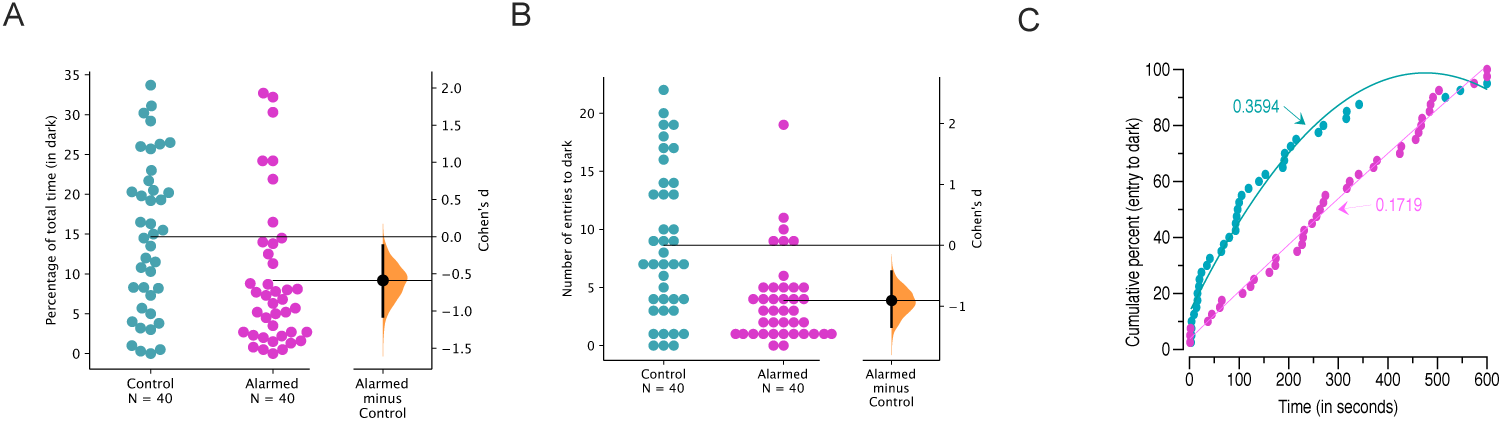
Effect of *Schreckstoff* on dark avoidance of larval zebrafish. A) The percentage of total time (of 10 minutes) spent in the dark, or B) the number of entries to the dark side and C) shows the time taken for cumulative percentage for the first entry to the dark side to reach 100% for control (teal) or alarmed (magenta) larvae. For A and B, Both groups are plotted on the left axes; the mean difference is plotted on floating axes on the right as a 5000 bootstrap sample distribution. The mean difference values are given in the text and are depicted as a dot; the 95% confidence interval is indicated by the ends of the vertical error bar.

### A lateral glomerulus in the olfactory bulb of larval zebrafish senses adult-derived *Schreckstoff*

Next, we examined the neural activity in the larvae. We first imaged the olfactory bulbs of 5-7 day old *Tg(gng8:GAL4, UAS:GCaMP6s)* larvae where a small subset of olfactory microvillus sensory neurons express GCaMP6s [41], and delivered *Schreckstoff* as described previously [14, 37]. A response to *Schreckstoff* was observed in the glomerular termini of gng8-expressing neurons (Figure 3 A-C). In the zebrafish, the position of large glomeruli is invariant as they are located in the same relative organization across individuals [8]. The developmental patterns of such large glomeruli can be traced from 72 hpf or 3 dpf onwards and reliably mapped even if the smaller glomeruli change in number and location in an experience dependent manner [9]. These are identifiable as in addition to the anatomical location in the X, Y and Z planes, the glomerular map can also be consistently derived on the basis of immunoreactivity to markers like calretinin and *Gαs/olf* [9].

**Figure 3.**
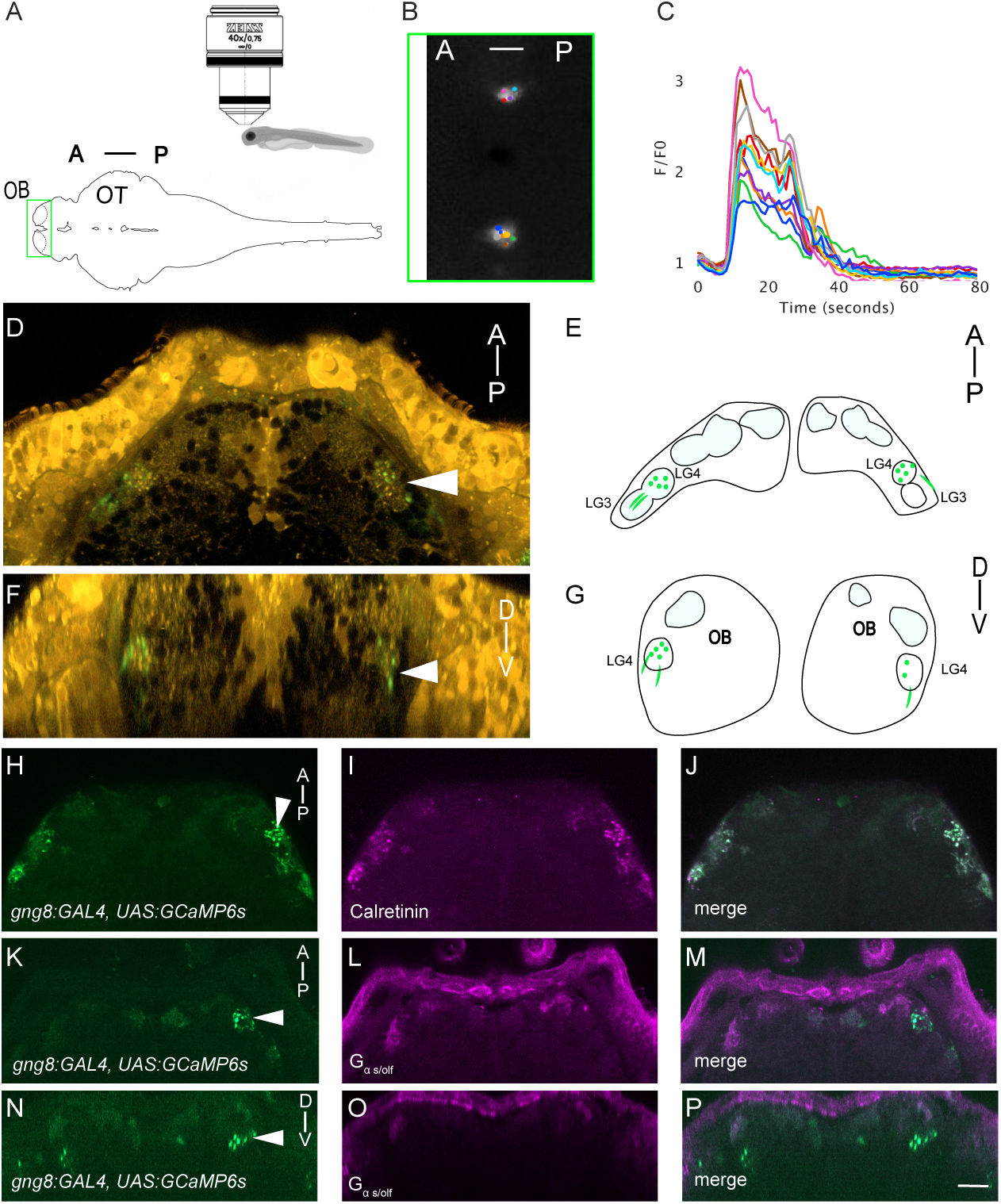
Larval OSNs targeting a lateral glomerulus respond to *Schreckstoff*. A) Schematic of the imaging setup for calcium imaging B) One frame from a F/F0 series, where fluorescence increase was maximal. C) Plot of 11 terminals, showing a rise in intracellular calcium in response to adult skin extract, which was delivered from 5 - 25 seconds. *Tg(gng8:GAL4, UAS:GCaMP6f)* fish, labeled with 4 − *Di* − 2 − *ASP* (shown in orange), which enables visualization of glomeruli. D) Dorsal view F) Frontal view through the stack. E) and G) show lateral glomerulus schematic based on D and F respectively. Labelled neurons (green) terminate in a lateral glomerulus. H) *Tg(gng8:GAL4, UAS:GCaMP6f)* stained with I) anti-calretinin and J) merge shows that transgenic neurons are Calretinin positive. K and N) *Tg(gng8:GAL4, UAS:GCaMP6f)* larvae stained with anti-*Gαs/olf* antibody visualized in L) dorsal and O) frontal views show no overlap in the O), and P) merge. A - Anterior and D - Dorsal is to the top in all images, P - Posterior, V - Ventral. Scale bar = 10 *µ*m.

We stained *Tg(gng8:GAL4, UAS:GCaMP6s)* larvae (Figure 3D-G) with anti-calretinin (Fig 3H-J) and anti-*Gαs/olf* (Figure 3K-P) antibodies. Antibody staining show that the *Schreckstoff* responsive glomerulus is calretinin positive but is not labeled by the anti-*Gαs/olf* antibody. The LG cluster consists of 2 identifiable large glomeruli, *LG*_3_ and *LG*_4_ from 3 dpf onwards in the classification proposed by Braubach et. al [9]. Among them, *LG*_4_ was reported to be the only glomerulus in the LG that is unresponsive to amino acids [9]. Therefore, based on the nomenclature proposed in [8], the *Schreckstoff* responsive glomerulus is located in the lateral glomerular cluster, most likely *LG*_4_, and is innervated by GnG8 positive neurons.

### Brain-wide responses to adult-derived *Schreckstoff*

Neurons from the larval olfactory bulb project to several regions in the forebrain, including the posterior telencephalon (Dp), ventral telencephalon (Vi and Vv) and dorsal right habenula [39]. To determine whether these regions show a change in activity as a result of exposure to the alarm substance, we recorded from the forebrains of 5-7 day old fish with broad expression of nuclear-localized GCaMP6f (*(Tg(elavl3:h2b-GCaMP6f)*), using a two-photon resonant scanning microscopy with piezo focusing to allow volume imaging (Figure 4 A, B). Correlated change in the activity was seen in the olfactory epithelium, olfactory bulb, Dp, Vi and habenula in all fish imaged (Figure 4 C-H; n = 7 fish). In addition to this correlated activity, we also observed a persistent increase in the activity in a subset of neurons in the lateral habenula of all fish (Figure 4 I, J; Supplementary Figure 6).

**Figure 4.**
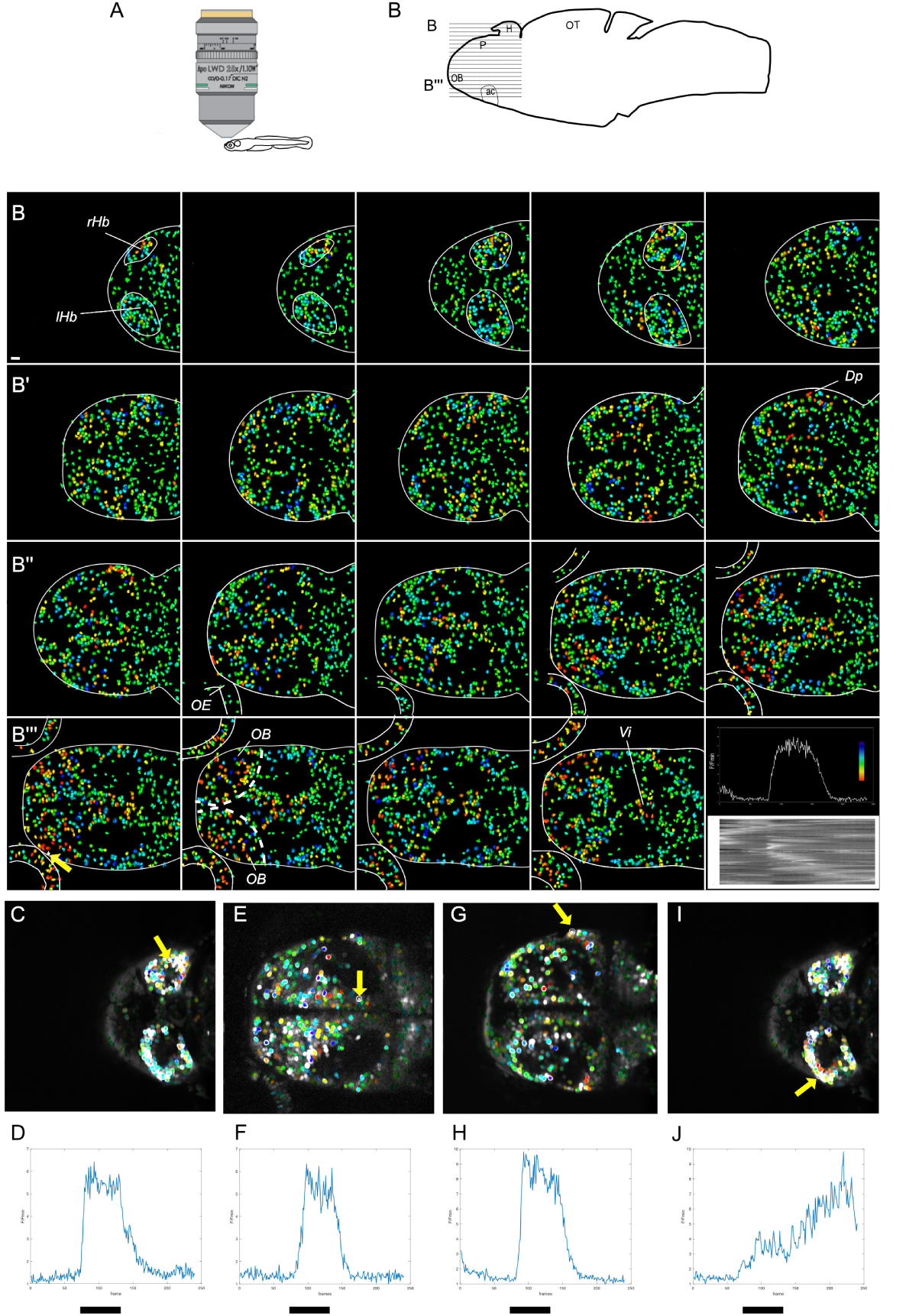
Forebrain responses to *Schreckstoff*. A) Schematic of the imaging setup for calcium imaging. B) Activity in nineteen focal planes (*B*′ to *B‴*) of the forebrain of a 7 day-old zebrafish, from dorsal (top left) to ventral (bottom row). Cells are color coded according to correlation with the indicated cell (bottom left panel), which is in the lateral olfactory bulb. The relative change in fluorescence of this cell is shown in the bottom right. The wedge shows the lookup table employed in mapping correlation. The raster plot shows raw fluorescence in > 7000 cells across all planes. C-J). Three different focal planes from the image in panel B, overlaid on a mean image of the fluorescence t-stack. These show three major targets of the olfactory bulb, which are Dp (C), Vv (E) and the habenula (G, I). Relative fluorescence change of the cell indicated indicated by the yellow arrow is shown in the plot below the corresponding image. The black bars indicate the stimulus. rHb - right habenula, lHb - left habenula, Dp - posterior telencephalon, Vi - intermediate ventral telencephalic nucleus, OE - olfactory epithelium, OB - olfactory bulb.

The lateral habenula receives direct input from the entopeduncular nucleus [2, 47], which is homologous to the internal segment of the globus pallidus in mammals. Imaging of larval fish expressing a fluorescent label in the entopeduncular nucleus *Tg(elavl3:h2b-GCaMP6f),etSqKR11* indicates that *Schreckstoff* elicits activity in this nucleus (Supplementary Figure 7). A mixture of activity patterns was seen, including activity during and after the stimulus (Supplementary Figure 6). Thus, in addition to areas predicted by direct connectivity with the olfactory bulb, the alarm substance influences a number of other areas within the forebrain, including a structure involved in processing negatively valenced stimuli [30].

The habenula provides a pathway from the forebrain to midbrain neurons. To determine if the habenular activity is accompanied by a change in midbrain activity, calcium imaging was carried out across a larger region of the brain (Figure 5A, n = 6 fish). Activity after exposure to the stimulus was distinct from activity before exposure (Figure 5B), indicating a change in brain state. In contrast to the olfactory bulb and telencephalon (Figure 5C-E), where there was a transient increase in activity that correlated with delivery of *Schreckstoff*, neurons in the midbrain tegmentum, superior raphe, posterior tuberculum and locus coeruleus showed persistent activity Figure (5F-H). These regions were identified based on anatomical landmarks as described in the methods. Thus, transient exposure to skin extract elicits an extended neuronal response in the diencephalon and midbrain.

**Figure 5.**
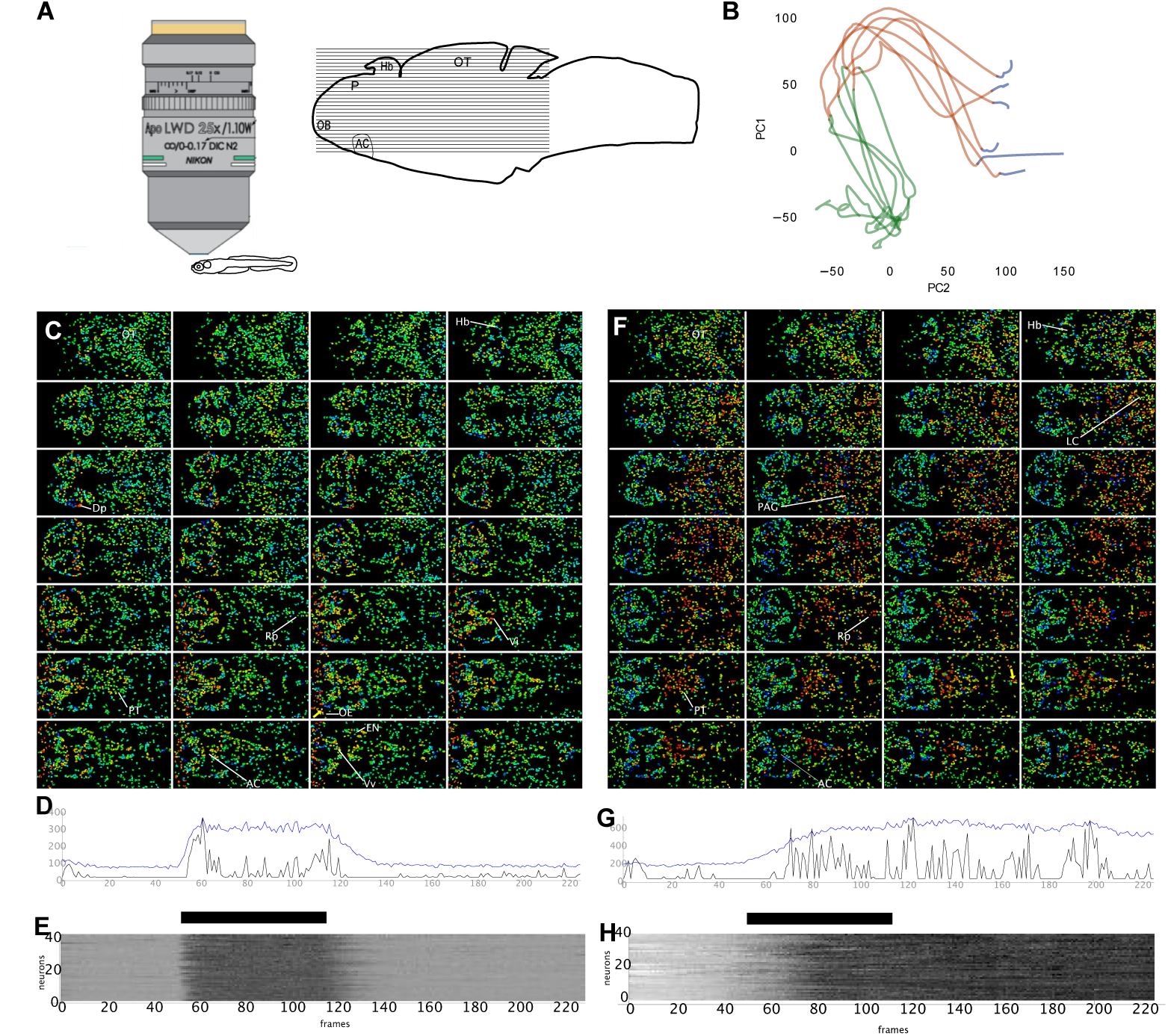
Contrasting dynamics in the forebrain and midbrain following*Schreckstoff* exposure. A) Schematic diagram of imaging set-up. The horizontal lines indicate the approximate location of the region imaged. 28 planes were recorded, 5 *µ*m apart. B) Trajectory of neural activity across the forebrain and midbrain of six fish, shown in two-dimensional state space. The colours indicate activity before (blue), during (red) and after (green) *Schreckstoff* delivery. C - H) Response in one 7 dpf fish to a pulse of *Schreckstoff*. Cells are colour-coded according to correlation coefficient with the indicated cell (yellow arrow), which is in the olfactory epithelium in panel C and in the raphe in panel F. D), G) The raw fluorescence (blue) and deconvolved trace (black) of the cell in panels C and F respectively. E), H). Heatmap of response in 40 cells that are most correlated with the cell indicated in panels C and F respectively. OT: optic tectum, AC: anterior commissure, Hb: habenula, Dp: Posterior telencephalon, Rp: superior raphe, LC: locus coeruleus, PAG: periaqueductal gray, EN: entopeduncular nucleus, Vi: intermediate ventral telencephalic nucleus.

## Discussion

In this study, we used the alarm response in larval zebrafish to characterize the change in brain state during danger. Two lines of evidence indicate that this is a valid paradigm. Firstly, behavioral studies, which were conducted in a size proportionate arena at an age when free-swimming and hunting begins, show a quantifiable change in locomotion in the presence of Schreckstoff. Following exposure, larvae show increased dark avoidance, suggesting that they are in a state of stress. This is consistent with previous studies showing an increase in cortisol levels in larval fish following exposure to Schreckstoff [20]. Secondly, calcium imaging indicates a reproducible change in global patterns of neural activity upon exposure to the alarm substance. Thus, the alarm substance is perceived as a threat by larval zebrafish, and induces a change in brain state.

A number of regions in the midbrain show sustained activity following exposure of larvae to the alarm substance. Activity in the superior raphe may lead to increased serotonin release, which has been shown to occur following exposure to the alarm substance in adult zebrafish [38]. Activity in the locus coeruleus is consistent with increased norepinephrine reported in adults following exposure to *Schreckstoff* [38] and is an evolutionarily conserved stress response of vertebrates. Sustained activity in posterior tuberculum, which contains dopaminergic neurons [17, 49], may similarly be part of a stress response [25, 26]. In the midbrain tegmentum, sustained activity was detected in a region that is proposed to be the periaqueductal gray, based on the expression of relaxin3a and proenkephalin-like [18]. This region is likely to regulate locomotion. We suggest that these are a part of the network that defines a stressed state, which remains even after a threat is removed.

These regions are likely to be regulated by the lateral habenula, which projects to the raphe, posterior tuberculum, and locus coeruleus. Lesioning of the lateral habenula blocks the response to *Schreckstoff* in adults [40], underscoring its role as a hub in the response network. The lateral habenula in turn, is known to receive input from the entopeduncular nucleus [2, 47]. This structure was active during and after stimulus delivery and receives input from the telencephalon [21, 28].

To identify circuits that could trigger the switch to a stressed state, we analyzed calcium imaging data for neurons that show a transient response that is correlated to delivery of the alarm substance, and which are anatomically upstream of neurons that show sustained activity. The alarm substance is detected by the olfactory system, and we observed responses in olfactory sensory neurons innervating a lateral glomerulus. This is likely to be LG4, which is the only glomerulus in this region that is unresponsive to amino acids [9]. Responses in the lateral olfactory bulb were correlated with activity in the lateral region of the pallium and in the ventral telencephalon. The former likely includes Dp (homolog of the olfactory cortex [23]) and Dlv (homolog of the hippocampus [23]), while the latter likely includes the intermediate nucleus of the ventral telencephalon, which receives input from the olfactory bulb and has been proposed to be homologous to the medial amygdala of mouse [5]. Based on the correlated activity and connectivity, we suggest that these telencephalic neurons contribute to the trigger circuit.

The mechanisms that sustain persistent activity in midbrain nuclei in the absence of an external trigger, and thus maintain the stressed state, remain unclear. Given the regulation of these nuclei by the lateral habenula and entopeduncular nucleus, it is likely that these are control points. The raphe itself is a potential source of activity in the entopeduncular nucleus [2], via a feed-forward loop, as are changes in excitability in the habenula [3]. Further investigation using the alarm response paradigm, combining behavioral studies with whole-brain imaging, connectomics and manipulation, should provide a greater understanding of the neural mechanisms underlying response to threat, and thus into conditions such as anxiety.

## Conclusion

Larval zebrafish can sense *Schreckstoff*. Most larvae (approximately 50%) show an immediate change in swimming behavior. Whole-brain imaging reveals a change in the brain state upon the perception of the alarm cue in the form of sustained activity in many mid-brain neuromodulator releasing regions. The larvae become vigilant after a brief exposure to the threatening cue, and transition to displaying defensive behaviors for an extended period even after the cue indicating danger is removed. Our study suggests that defensive behaviors operate over a continuum and involve multiple brain circuits.

## Financial Disclosures and Acknowledgement

This research was supported by a Lee Kong Chian School of Medicine, Nanyang Technological University Singapore Start-Up Grant and by the Singapore Ministry of Education under its Academic Research Fund Tier 2 Award (MOE2017-T2-058) to SJ, and by Yale-NUS College grants R-607-265-225-121 and IG16-LR003 to ASM. We thank Caroline Kibat for assistance with immunostaining, and staff at Zebrafish Fish Facility, IMCB.

## Author contribution

SJ - Conceptualization; Data curation; Formal analysis; Funding acquisition; Investigation; Methodology; Resources; Visualization; Writing - original draft; Writing - review & editing. SK - Data curation; Formal analysis; Investigation; Methodology; Writing - review & editing RKC - Data curation; Formal analysis; Investigation; Methodology; Writing - review & editing ASM - Conceptualization; Data curation; Funding acquisition; Investigation; Methodology; Resources; Visualization; Project administration; Writing - original draft; Writing - review & editing.

