## Supplementary material and methods for "Neural correlates of state transitions elicited by a chemosensory danger cue"

#### Supporting Information

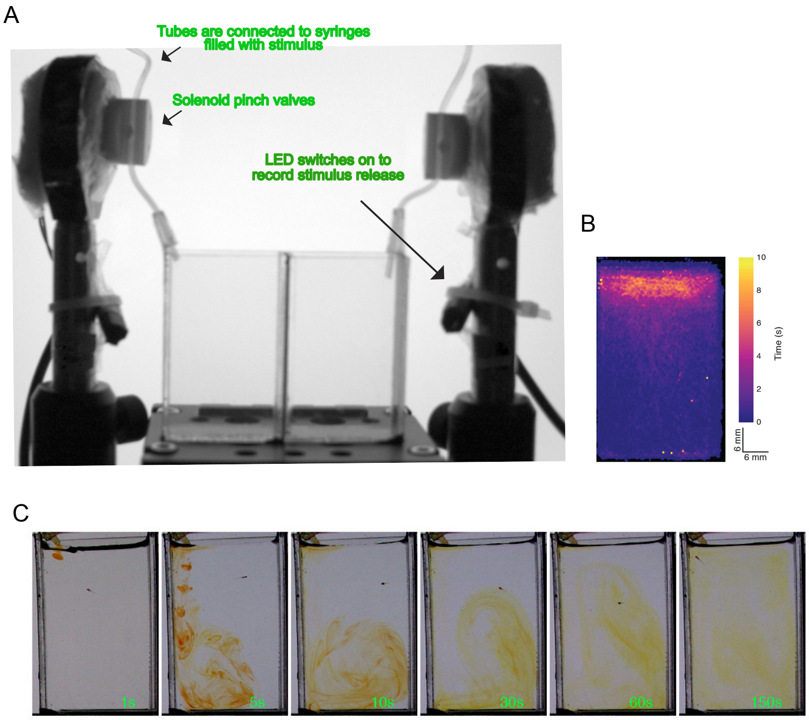


**S1 Fig.**
**Experimental setup for olfactory experiments.** (A) Fish were tested in pairs in the setup shown. Stimulus delivery was performed via the tubes whose opening was controlled by solenoid pinch valves actuated by a 12V DC input. LED switches (not visible to fish) were used to identify the delivery time in the videos acquired. (B) Representative heat map showing the spatial location of a fish in 30 min recordings. Fish normally preferred swimming in the top third of the tank. (C) Dispersal of a Phenol-red dye delivered for the same duration as the stimulus (5 seconds) in the tank over time (shown in the lower corner). The larval fish is visible as a dark spot.

#### S2 Fig.

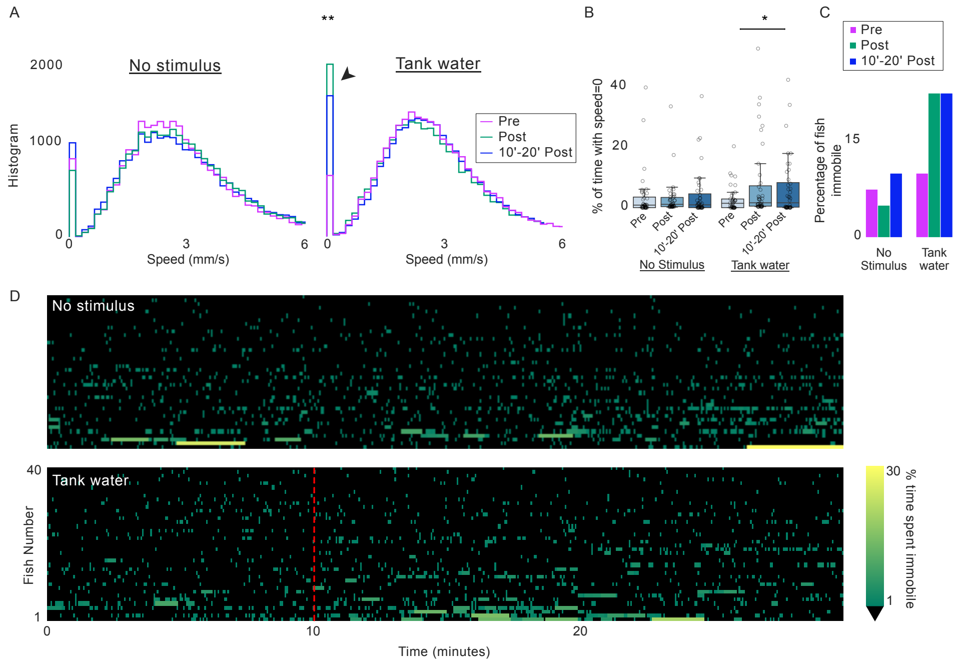


**Stimulus delivery startles larval zebrafish** (A) Histograms of speed distribution (in mm/s) to A) no stimulus and control stimulus (tank water) delivery. The distribution is derived from 3 ten-minute bins before (Pre), ten minutes after (Post) and ten to twenty minutes after (10'-20' Post) control stimulus (tank water) delivery. For comparison, the no-stimulus condition data was also divided into the same bins. No significant difference in Pre with Post: p=0.3599; Pre with 10'-20' Post: p=0.2184 (KS test) in no stimulus condition, but stimulus delivery changes were significant Pre with Post: p=0.0017; Pre with 10'-20' Post: p=0.0053 (KS test). Increase in time with speed = 0 mm/sec after stimulus delivery (arrowhead). (B) Boxplots of the percentage of time spent motionless in the three time-bins for the two conditions. Circles represent the individual fish response. Pre with Post: p=0.025 and Pre with 10'-20' Post: p = 0.030, Student's t-test (C) Percentage of fish in the two conditions (n = 40 per condition) that show a speed = 0 mm/sec for 1 SD longer than the average immobility time in the Pre time-bin of the no stimulus condition. Approximately 20% of the fish show a response when a stimulus is delivered. (D) Heatmaps show the start and the duration of events with speed = 0 in the two conditions. In the color scale, black represents speed > 0 mm/sec or periods of activity. * indicate p values < 0.05, and ** < 0.01, and *** indicate < 0.0001 in paired or unpaired Student's t or KS test. Exact p values are given in the text.

#### S3 Fig.

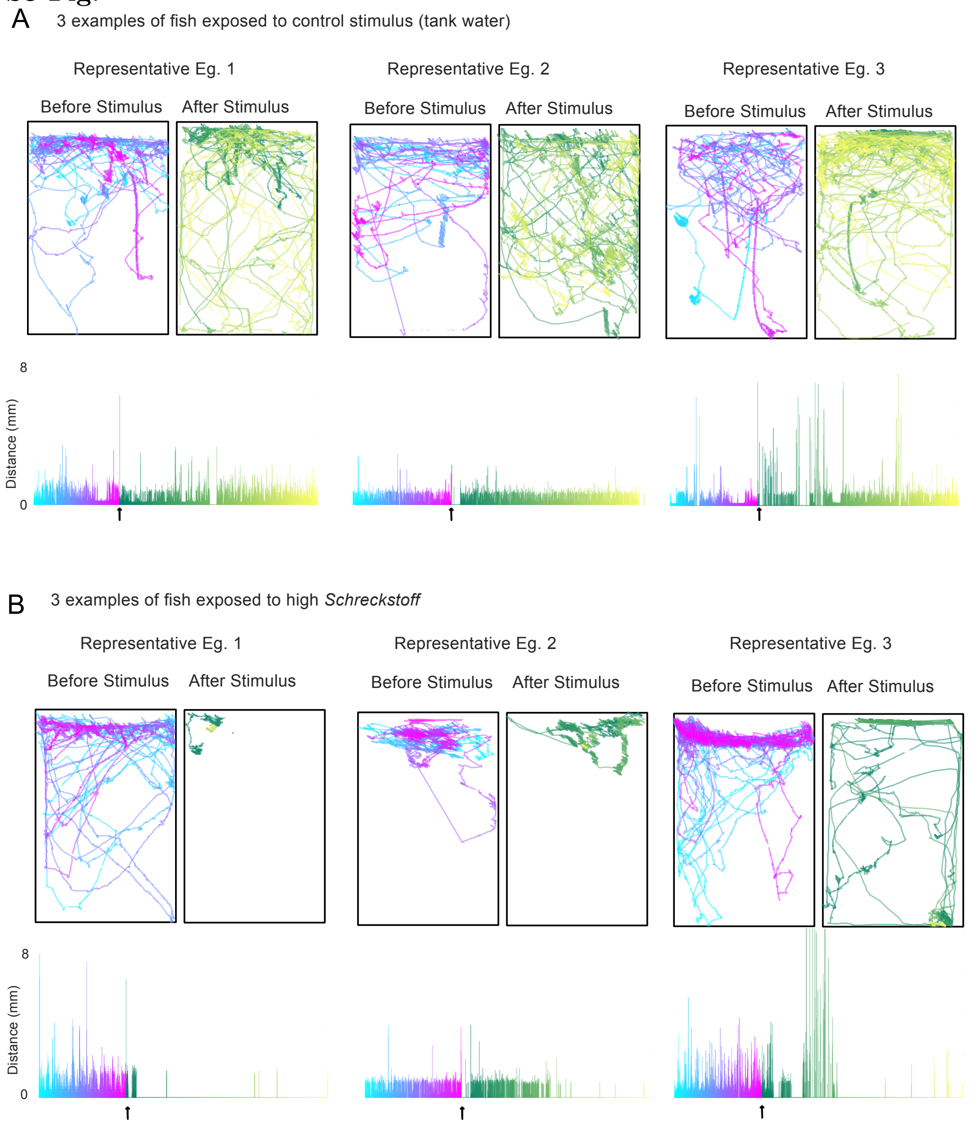


**Representative traces from 3 larvae** Tracks and distance swam (in mm) color-coded over the Pre and Post periods exposed to A) Control (tank water) or to B) a high concentration of *Schreckstoff*.Stimulus delivery is indicated by arrows.

#### S4 Fig.

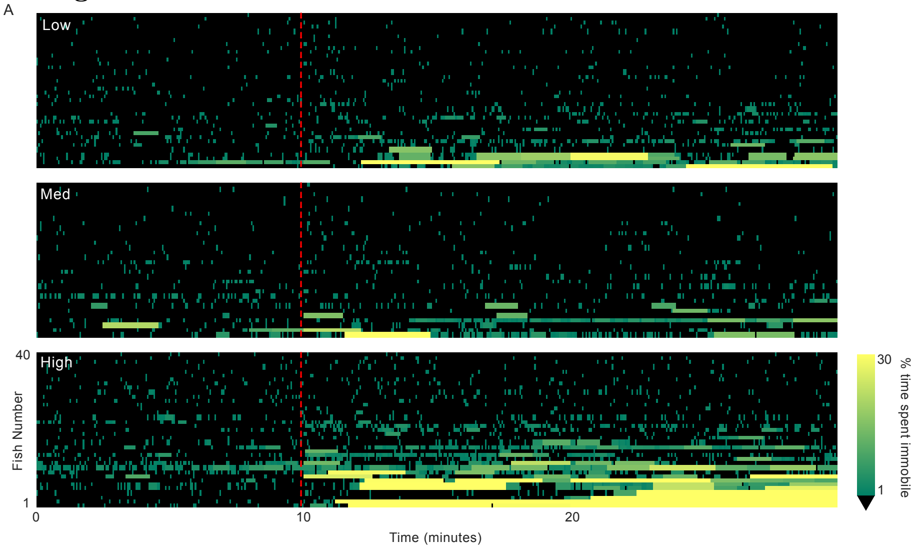


**Larvae show an alarm response to high concentrations of *Schreckstoff***
(A) Heatmaps show the start and the duration of events with speed = 0 in the two conditions. In the color scale, black represents speed >0 mm/sec or periods of activity.

**S5 Fig.**

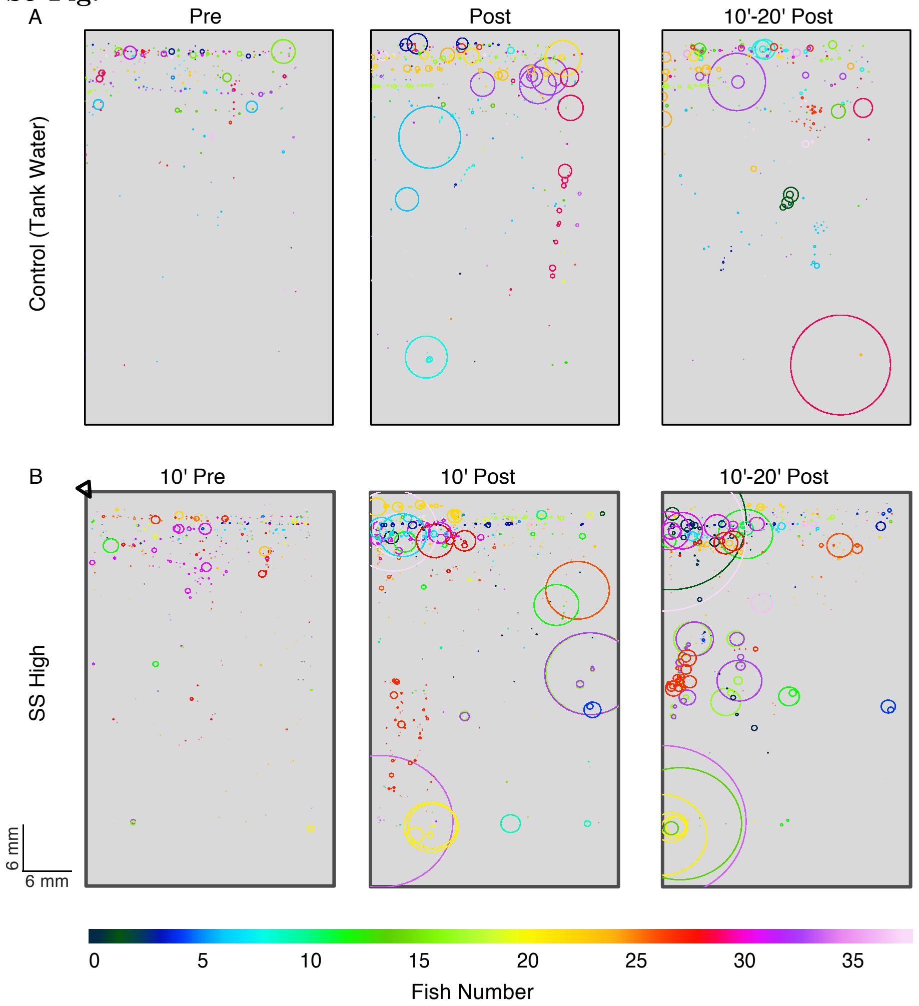


**Larvae do not show a diving response to *Schreckstoff*** Spatial representation of areas in the tank where the speed was = 0 mm/sec in A) control and B) High *Schreckstoff* conditions. Circles were drawn on the centroid of the fish. The size of the circle indicates the length of time the fish stayed in that position. Each fish in the experiment is represented by a color in the color bar. The location of the stimulus release is indicated by the black triangle in B.

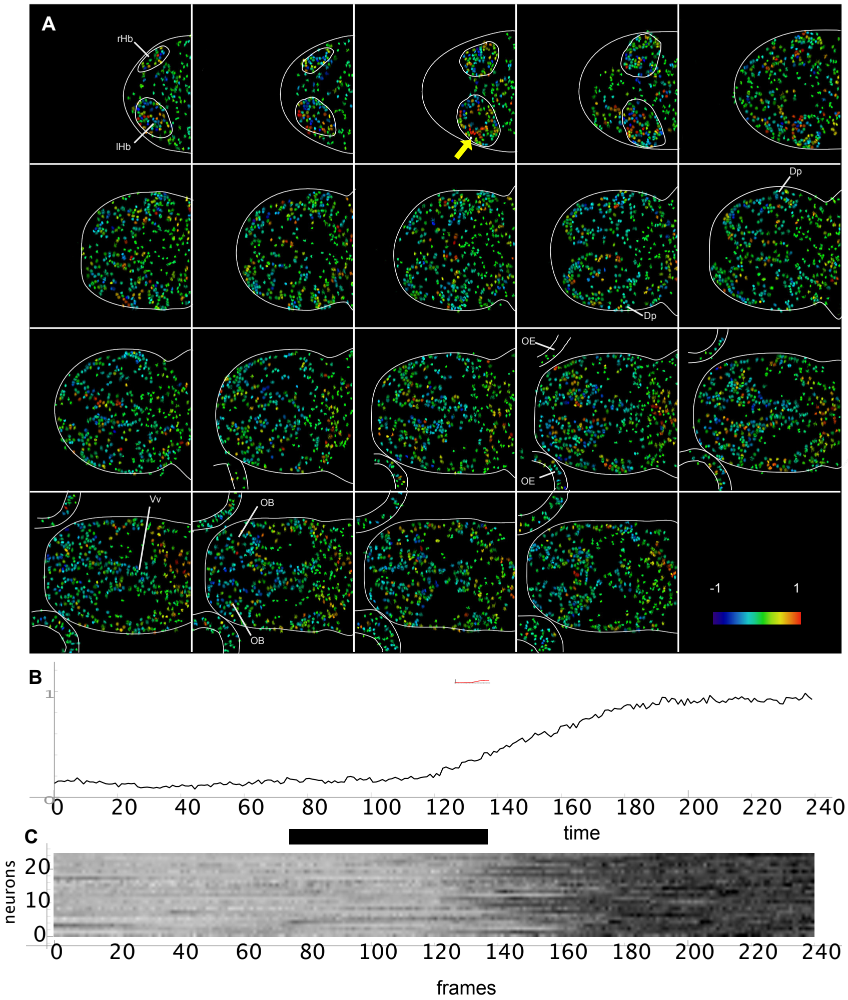


**S6 Fig.**

**Sustained forebrain response to *Schreckstoff*.** (A) Neural activity in the forebrain in response to a pulse of Schreckstoff. This is the same recording as shown in Figure 6, except that neurons are colour coded according to correlation coefficient with a cell in the lateral habenula (yellow arrow). (B) The average change in activity in 25 cells that are most highly correlated with the indicated cell. The bar indicates the time when the olfactory neurons showed a response. (C) Heatmap showing activity in the 25 cells that are most correlated with the indicated cell. rHb: right habenula, lHb: left habenula, OB: olfactory bulb, OE: olfactory epithelium, Dp: posterior telencephalon, Vi: intermediate ventral telencephalic nucleus.

**S7 Fig.**

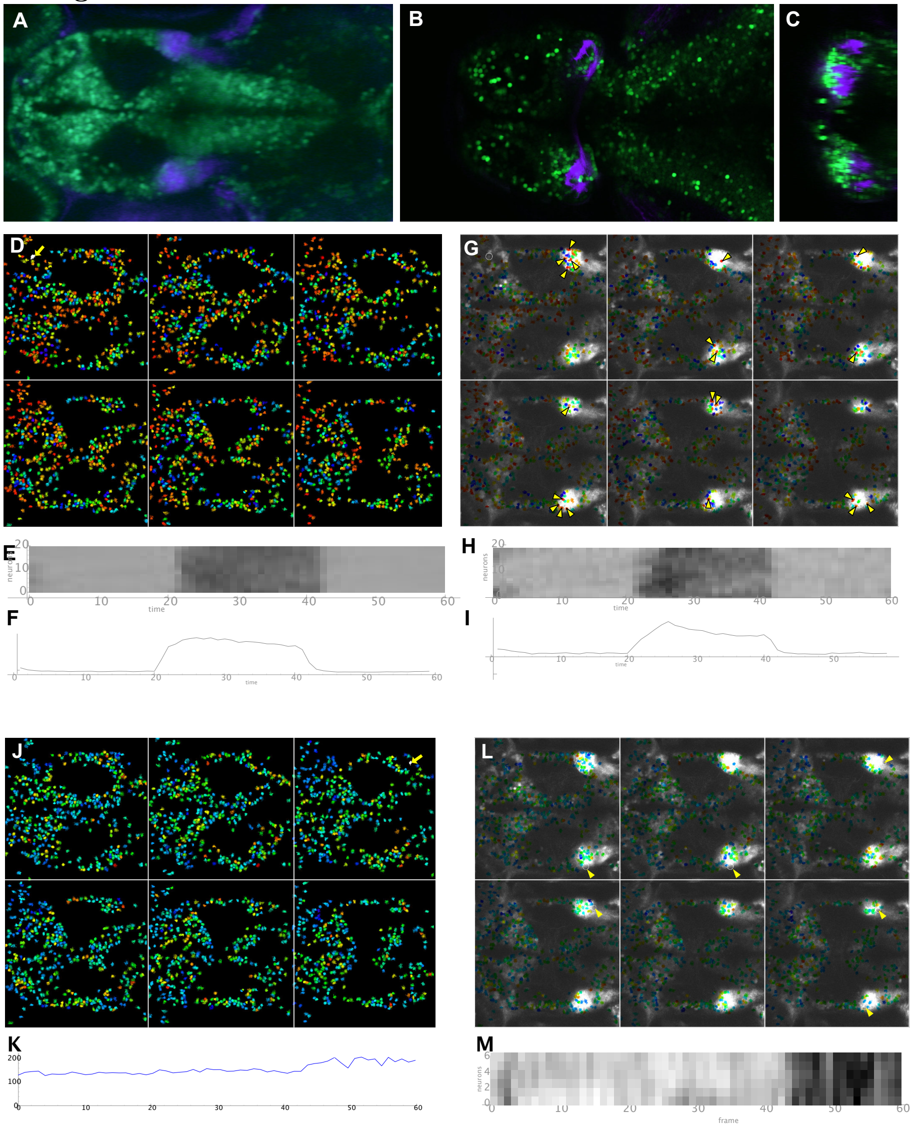


**Activity in the entopeduncular nucleus.** (A-C) Dorsal view of a larval fish with expression of a fluorescent reporter in neurons located within the entopeduncular nucleus (A), which extend axons to the habenula (B,C). (C) is a transverse section through B, at the position indicated. (D-I) Neural activity in six different focal planes, 5 $\mu$m apart, in the forebrain, in response to transient exposure to Schreckstoff. (D) Neural response, correlated to a cell in the dorso-lateral olfactory bulb (yellow arrow). E. Heat-map showing the response of twenty cells that are most highly correlated. (F) Average response of the twenty cells in E. G. The activity map from panel D, overlaid on an image showing the entopeduncular nucleus labelled in the EtKR11 line. (H) Response of 20 cells in the entopeduncular nucleus (yellow arrowheads), which are correlated with the cell in the olfactory bulb (circled in panel (G). (I) Average of the entopeduncular excitatory response shown in panel H. (J-M) Post-stimulus activity in the entopeduncular nucleus. (J) Neural activity, correlated to the cell indicated by the yellow arrow. K. Raw fluorescence of the cell indicated in panel J. (L) The activity image from panel J, overlaid on an image showing the entopedencular nucleus labeled by EtKR11line. (M) Heat map of 6 cells in the entopeduncular nucleus indicated in panel L. There is an increase in fluorescence after delivery of the stimulus.

**S1 Video. Example video showing alarm response in 5-7 day old larva**.

### Extended Methods

#### Preparation of alarm substance from adult zebrafish

Alarm substance or *Schreckstoff* was prepared as described previously [7]. Briefly, 3-4-month-old adult zebrafish were euthanized by immersion in ice-cold water for 10 minutes. Fish were dried on KimWipes and 7 to 8 shallow lesions were made using a microsurgical knife. Fish were immersed in an Eppendorf with 2ml of tank water and shaken gently on an orbital shaker for a minute. Fish were removed and the crude extract was then heated at 95°C overnight (  8 to 10 hours). The extract was then centrifuged at 13.2k rpm and filtered. The crude extract was aliquoted into different concentrations for the larval experiments. The highest concentration was 1:500 dilution of the crude extract. The medium and low concentrations were 1:10 and 1:100 of the highest concentration respectively.

#### Olfactory behavior assay procedure

All experiments were performed between 1000-1700 hrs. Tanks were sanitized with alcohol at the end of each experiment and thoroughly washed with the system or tank water. Silicon tubing was replaced when testing different concentrations of stimuli to avoid contamination. Fish were acclimated in the vertical tanks for 5 minutes before being placed in the experimental chamber. Only larvae that started swimming and exploring the tanks within 5 minutes were used in the experiments (approximately 1 in 20 larvae were rejected). Once in the setup, Pre (10'), Post (10') and 10'-20 Post (10') were recorded with stimulus delivery triggered as described below.

#### Olfactory behavior assay

The behavior setup and protocol are shown in Supplementary Fig S1 Fig. The entire setup was placed in a controlled environment (incubator) to remove external light, sound, or vibrations. A pair of larvae were tested at a time in two tanks with dimensions 30 mm x 10 mm x 50 mm (LxWxH). Larvae in this chamber can swim across the length and height of the observation chamber in approximately 4 to 6 body lengths but have restricted movement in the width to about 1.5 body lengths. The tanks were filled with system water that was also used as a control stimulus. The sides of the tank were covered to obscure fish from each other. Stimuli were delivered through two silicon tubes (Automate Scientifica). Delivery was controlled by solenoid actuation via a TTL pulse delivered by an Arduino Uno board. A USB 3.0 Basler acA2040-90umNIR was used for acquiring high-speed videos (60 fps) placed 15 cm in front of the tanks. A white light LED lightpad (Artograph) was used to illuminate tanks to provide a constant and uniform backlight. Custom written Python codes controlled video acquisition and triggering the TTL pulse. Videos collected were tracked offline by custom-written Python scripts as described below.

#### Olfactory behavior locomotion analysis and statistics

Larvae were tracked using custom-written Python codes (available at $https://github.com/seethakris/zebrafish_tracking$). The algorithm tracked pixels that changed between consecutive frames. The output of tracking is the coordinates of the centroid of the fish across time. About 12.5% of the videos were excluded because the tracking algorithm could not accurately track a significant portion of the video (120 seconds or more). This happened if the fish were obscured on the edge of the tank or were at the meniscus. Boxplots are used to show distribution of data points. In the boxplots, the box ranges from the first quartile to the third quartile of the data and shows the interquartile range (IQR). The line across the box is the median, and the whiskers extend to 1.5 * IQR on either side of the box. Anything above this range is an outlier and is shown by a black diamond. Distribution of speed in fish in all conditions were normal distributions (p¡0.0001, Shapiro-Wilk test) and thus, reported p-values and test statistics were calculated using parametric paired or unpaired t-test. Effect sizes were calculated using Cohen's *d*.

#### Light/dark choice assay

All experiments were performed between 1000 − 1700 hrs in a behavioral arena within blackout curtains as described previously [2]. Four transparent plastic tanks (dimensions: 43 mm W × 60 mm L × 30 mm H) filled with 30 mL fresh tank water served as assay chambers. Opaque cardboard sheets were placed between the 4 tanks. Tanks were placed on an Apple iPad screen and videos (9fps) were recorded on a USB3.0 Basler camera (Model# acA2040-90umNIR 1440 × 1080 pixels) with a long-pass filter (MIDOPT, LP830 830nm) placed above. Two IR light bars (850 nm peak from TMS-lite) positioned next to the 4 tanks served as an IR light source. The light/dark compartments were created by white and black rectangles (with 50% transparency level for the black) on a Microsoft PowerPoint slide displayed on the iPad Air at highest brightness level. Larvae were exposed either to *Screckstoff* or to control (tank water) in a treatment tank of the same dimensions with the same volume of tank water for 10 minutes. Larvae were then gently pipetted from the treatment tank into two wash-out tanks sequentially before being placed into the dark/light test chamber. Real-time tracking of the fish was achieved by custom-written Python codes using the OpenCV library and recorded as coordinates of the centroid of the fish across time. Larval behavior was recorded for 10 minutes. Offline analysis was performed using custom-written macros to determine the position of the animals, the number of entries into each compartment, the percentage of total time spent in each compartment, etc. A few fish (n=1 in the 7dpf Control group and n=1 in the 13dpf Alarmed group) were excluded from the analysis because they were immobile or because of tracking errors.

#### Calcium imaging

Larvae were immobilized using Mivacurium (Tocris), then mounted in 2% low melting temperature agarose on a rectangular glass coverslip. A wedge of agarose was removed from the front of the fish. Odor was delivered using a perfusion device (Warner VC-6M), with the aid of a RC-26GLP open diamond bath mounted on the PM1 platform. Time-lapse imaging was carried out on a Nikon A1RMP microscope using a 20x water immersion objective (NA 1.1). Images were collected with no delay between frames, using the resonant scanner and no averaging. The focus was controlled with a piezo driver (Mad City Labs). Images were registered, segmented and analyzed using Suite2p [6].

#### Anatomy

Brain regions were identified on the basis of morphological landmarks and gene expression patterns. The periaqueductal gray (PAG) in vertebrates is defined as a midbrain region surrounding the aqueduct, which extends posteriorly from the tegmentum, in a region dorsal to the interpeduncular nucleus. In identifying the structure in larval zebrafish, these anatomical landmarks as well as the expression of relaxin3a and proenkephalin [3], and tachykinin [4] were used. A similar region has been defined in the lamprey [5]. The intermediate nucleus of the ventral telencephalon is defined as the region caudal and dorsal to the anterior commissure [1].
